# Mixed selectivity in the cerebellar Purkinje-cell response during visuomotor association learning

**DOI:** 10.1101/2021.08.12.456160

**Authors:** Naveen Sendhilnathan, Anna E. Ipata, Michael E. Goldberg

## Abstract

Although the cerebellum has been traditionally considered to be exclusively involved in motor control, recent anatomical and clinical studies show that it also has a role in reward-processing. However, the way in which the movement related and the reward related neural activity interact at the level of the cerebellar cortex and contribute towards learning is still unclear. Here, we studied the simple spike activity of Purkinje cells in the mid-lateral cerebellum when monkeys learned to associate a right or left-hand movement with one of two visual symbolic cues. These cells had distinctly different discharge patterns between an overtrained symbol-hand association and a novel symbol-hand association, responding in association with the movement of both hands, although the kinematics of the movement did not change between the two conditions. The activity change was not related to the pattern of the visual symbols, the movement kinematics, the monkeys’ reaction times or the novelty of the visual symbols. The simple spike activity changed with throughout the learning process, but the concurrent complex spikes did not instruct that change. Although these neurons also have reward-related activity, the reward-related and movement-related signals were independent. We suggest that this mixed-selectivity may facilitate the flexible learning of difficult reinforcement learning problems.

## Introduction

Historically, the cerebellum has been considered to be exclusively involved in the acquisition of motor skills (De Zeeuw and Ten Brinke, 2015), motor learning, and gain adaptation (Thach, 2012). However, recent converging evidence (Heffley and Hull, 2019; Heffley et al., 2018; Kostadinov et al., 2019; Larry et al., 2019; Sendhilnathan et al., 2019, 2020) strongly suggests that the cerebellum also processes reward related information. In particular, the lateral area of Lobule VII of the cerebellar hemisphere, near Crus I and II, is a region that has extensive reciprocal connections with other reward processing areas of the brain including prefrontal cortex and basal ganglia (Caligiore et al., 2017; Koziol et al., 2014; Tedesco et al., 2011). When monkeys learn to associate an arbitrary symbol with a well-learned movement in a reinforcement learning paradigm, Purkinje cell (P-cell) simple spikes produce an error signal, which decreases as the monkeys learn the task (Sendhilnathan et al., 2020). Transient inactivation of this area impaired the monkeys’ ability to learn a new set of associations (Sendhilnathan and Goldberg, 2020).

Although the cerebellum receives both motor related (Raymond and Medina, 2018) and reward related information (Wagner et al., 2017), their interaction at the level of cerebellar cortex and their contribution to learning is still unclear. To study the nature of information encoded by the cerebellum during learning an arbitrary stimulus-response association, we compared the activity of P-cells in Crus I and II while two monkeys were actively learning new visuomotor associations and while they performed an already well-learned, familiar visuomotor association task. We found that the simple spikes of hand-related P-cells changed their activity profile significantly when we changed the visuomotor association from a well-learned to a novel association, even though the monkeys made the precisely same hand movement to report their choices. The pattern of the simple spike response continued to change throughout the learning process. This change in neural activity was independent of the kinematics of the movement, the hand used to report the choice, the complexity of the visual symbols, the reaction time of the animal, or the novelty of the visual symbols. Furthermore this signal could be dissociated from the reward-related error signal, which described the result of the monkey’s prior decision (Sendhilnathan et al., 2020). The concurrent complex spike activity was unlikely to have instructed this change in simple spike activity. Our results provide evidence that P-cells in the lateral part of Lobule VII of the cerebellar hemispheres multiplex two different signals when monkeys learn a new arbitrary stimulus response association. One is the previously described reinforcement learning error signal (Sendhilnathan et al., 2020), and the second, described here is a signal that describes the state of learning. Neither participates in encoding specific motor kinematics. We suggest that this mixed selectivity facilitates the flexible learning of a new visuomotor associations (Rigotti et al., 2013).

## Results

Two monkeys performed a two-alternative forced-choice discrimination task, where the monkeys learned to associate one of two visual symbols with a left-hand movement and the other symbol with a right-hand movement. The monkeys began each trial by placing their hands on each of two manipulanda, after which one of the two symbols appeared on the screen and the monkeys lifted the hand that was arbitrarily associated with that symbol to earn a liquid reward (**Fig 1b**). We first trained the monkeys to associate a specific pair of symbols (green square and pink square) with specific hand movements (left and right-hand release, respectively) for about 4-6 months until their performance was above 95% correct; we refer to this as the overtrained condition (**Fig 1c**).

**Figure 1:**
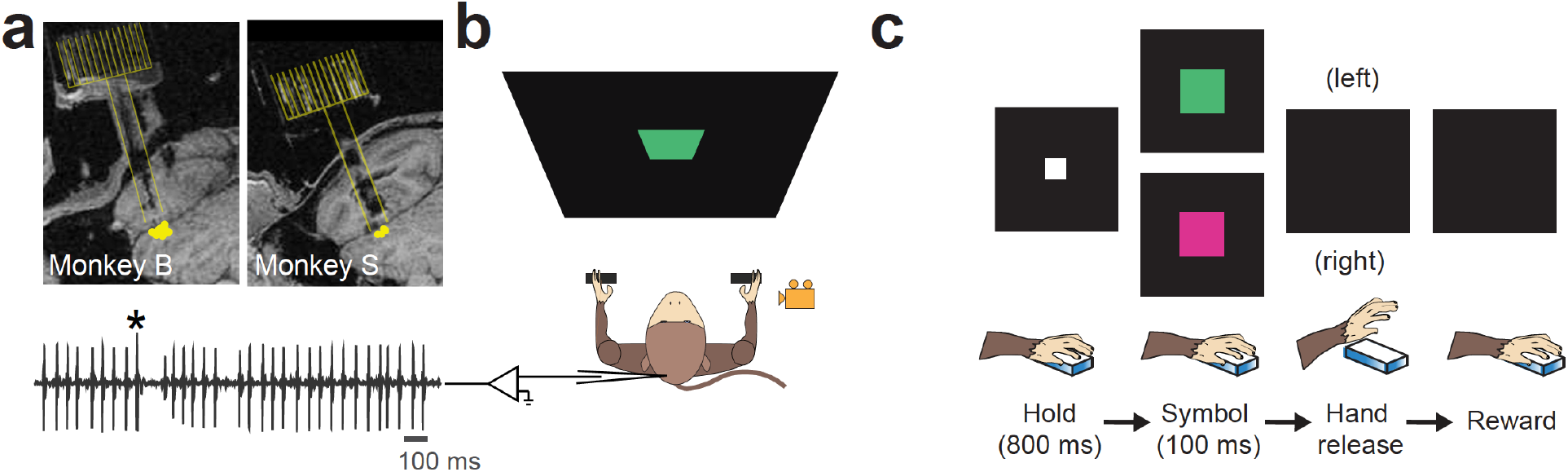
Visuomotor association learning paradigm. **a**. Top: T1-MRI with chamber location showing the recording locations in the mid-lateral cerebellum of two monkeys. Bottom: Raw neural recording showing simple spikes and a complex spike (marked by *). **b**. Schematic of the recording set up. The monkey was seated on a chair facing a screen where the symbols were back-projected. The monkey grabbed and released manipulanda with either hand during the task. The right-hand movement was recorded using a high-speed video camera (shown in yellow). **c**. Two-alternative forced-choice discrimination task (top) and parameters (bottom): The monkeys pressed each bar with a hand, and a white square appeared that served as the cue for the start of the trial. Then one of the two visual symbols appeared briefly. The monkey lifted the hand associated with the presented symbol to earn a reward immediate. The correct symbol-choice association is shown for the overtrained task (Green symbol – left hand; Pink symbol – right hand).

### Visuomotor association learning related changes in simple spike activity were independent of motor kinematics

We began each experiment with the overtrained symbols and then after ~30 trials, we switched the symbols to arbitrary fractal symbols which the monkey had never seen before. We refer to this as the novel learning condition. The monkeys performed the overtrained task with close 100% accuracy. However, once we switched the symbols to novel symbols, their performance dropped to chance level. The monkeys learned the arbitrarily assigned correct symbol-hand association through trial and error, usually in ~50-70 trials on an average, through an adaptive learning mechanism.

Here, we describe further analysis of the activity of single P-cells recorded in Lobule VII near crus I and II of the cerebellar hemisphere, while monkeys performed the visuomotor association task (Sendhilnathan et al., 2020). During the overtrained condition, most P-cells (106/128) significantly increased their firing rate during the bar-release hand movement. Therefore, given the prominent role of the cerebellum in motor control (Manto et al., 2012; Thach, 2012), if the P-cells only encoded the kinematics and the motor error of the hand movements, as long as the hand movement made by the monkeys remained the same, we would expect the neural activity to not change when the monkeys had to learn a new visuomotor association. However, contrary to this, we found that when the task changed from the overtrained to the novel condition (**Fig 2a**), the activity of 105/128 P-cells changed dramatically (*P* <0.001; KS test) in several intervals of the trial (for single cell example, see **Fig 2b**). Across the population, the change in activity occurred at different epochs for different neurons (for single cell examples, see **Fig S1)**.

**Figure 2:**
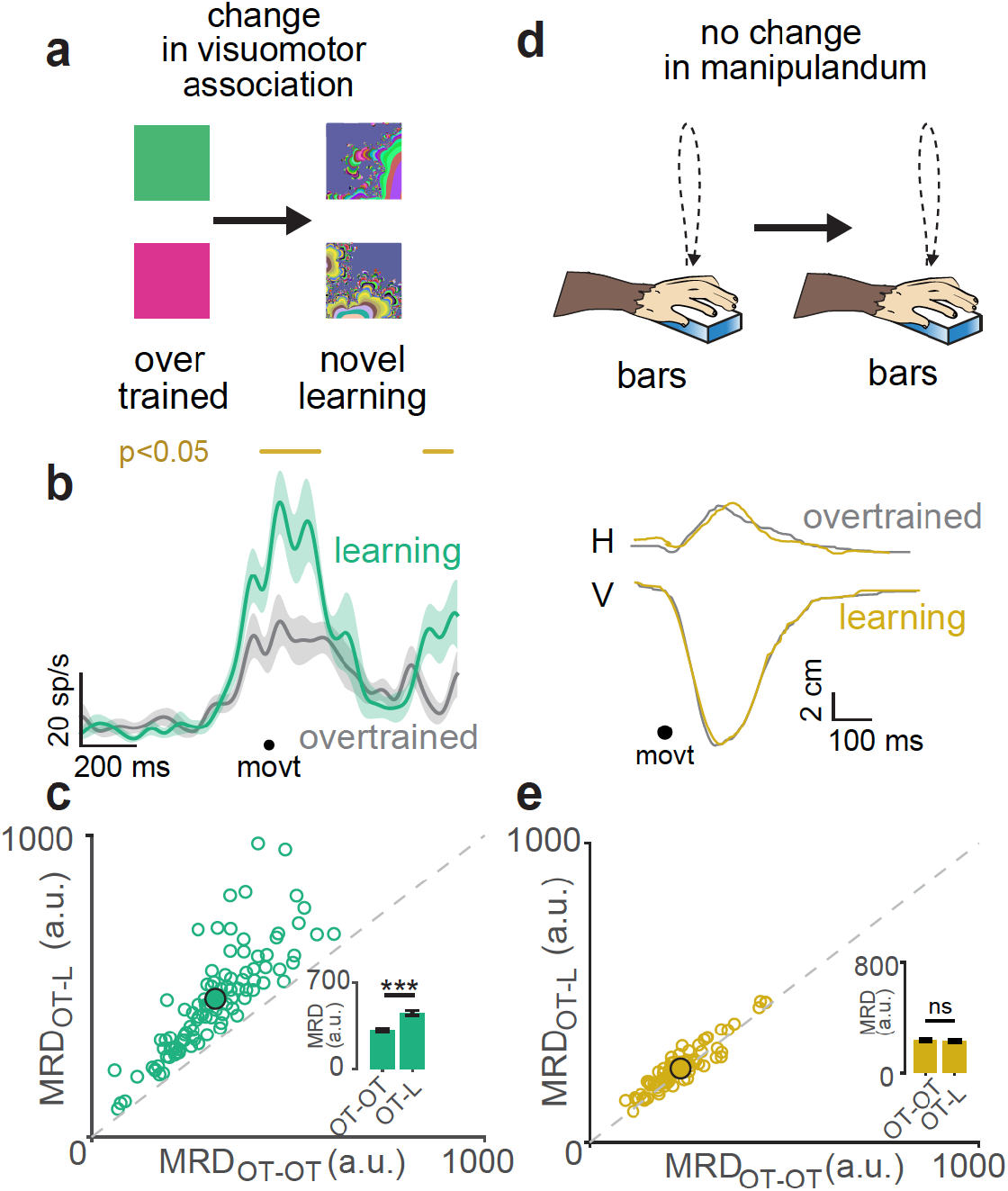
P-cell activity distinguished overlearned and novel symbols despite the absence of changes in movement kinematics. **a**. Task in which there was a change in visuomotor association from overtrained to novel association. **b**. Spike density plot of a representative P-cell aligned to bar-release hand movement onset (movt) that responded to the symbol switch by firing differently between the overtrained (gray) and novel (green) conditions. The epochs where this difference was significant (p<0.05, t-test) is shown by a gold line on the top. **c**. Scatter plot of Mean RMS Distance (MRD) for within overtrained (OT) condition (OT-OT) vs MRD for across overtrained and novel learning (L) condition (OT-L), obtained from method described in **Fig S2**. Each open circle is a P-cell and the mean value is shown as a filled circle. The inset shows the mean MRD for OT-N and OT-OT conditions. *** means *P* <0.001, Mann-Whitney U-test. **d**. A cartoon bar-release hand movement trajectories (top) and the actual movement trajectories decomposed into horizontal (H) and vertical (V) components traces (bottom) for overtrained (grey) and novel (yellow) conditions. **e**. Same as **c** but for bar-release hand movement trajectories. n.s means *P* = 0.3822, paired t-test

To show that the activity profile between the two conditions was significantly different despite this heterogeneity, in a way that is neither epoch nor neuron (statistical distribution) dependent, we computed the root mean squared (rms) distance between the spike density functions across the overtrained and novel conditions with repeated random sampling and compared the resulting distribution’s mean (mean rms distance, MRD_across_) with the mean from a distribution of two random samples repeatedly drawn without replacement, from within the overtrained condition (MRD_within_), compensating for differences due to reaction times (**Fig S2**; see methods). The within-condition MRD values, for the population, were significantly lower than the across-condition MRD values indicating that the change in neural activity in the novel condition was significantly different from the activity in the overtrained condition (*P* <0.001 Mann-Whitney U-test; **Fig 2c**). To see if the change in P-cell activity were accompanied by a change in the kinematics of the monkey’s movement, we painted a fluorescent dot on the monkey’s hand, and recorded its x-y position with time, using a video camera running at 200 frames/s (**Fig S3**; see methods). Although the neural activity changed dramatically at the symbol switch the monkeys showed no significant difference in motor kinematics between the overtrained and novel conditions (single session example: **Fig 2d** and population: **Fig. 2e**, *P* = 0.3822, paired t-test).

Furthermore, the change of neural activity at the symbol switch was unrelated to any change in reaction time at the symbol switch: Although in most experiments the reaction time increased at the symbol switch (**Fig 3a**), the reaction time did not change at the symbol switch on 24/105 (23%) sessions (**Fig 3b**) even though the monkeys’ performance decreased significantly and the neural activity changed significantly in these sessions (**Fig 3c**).

**Figure 3:**
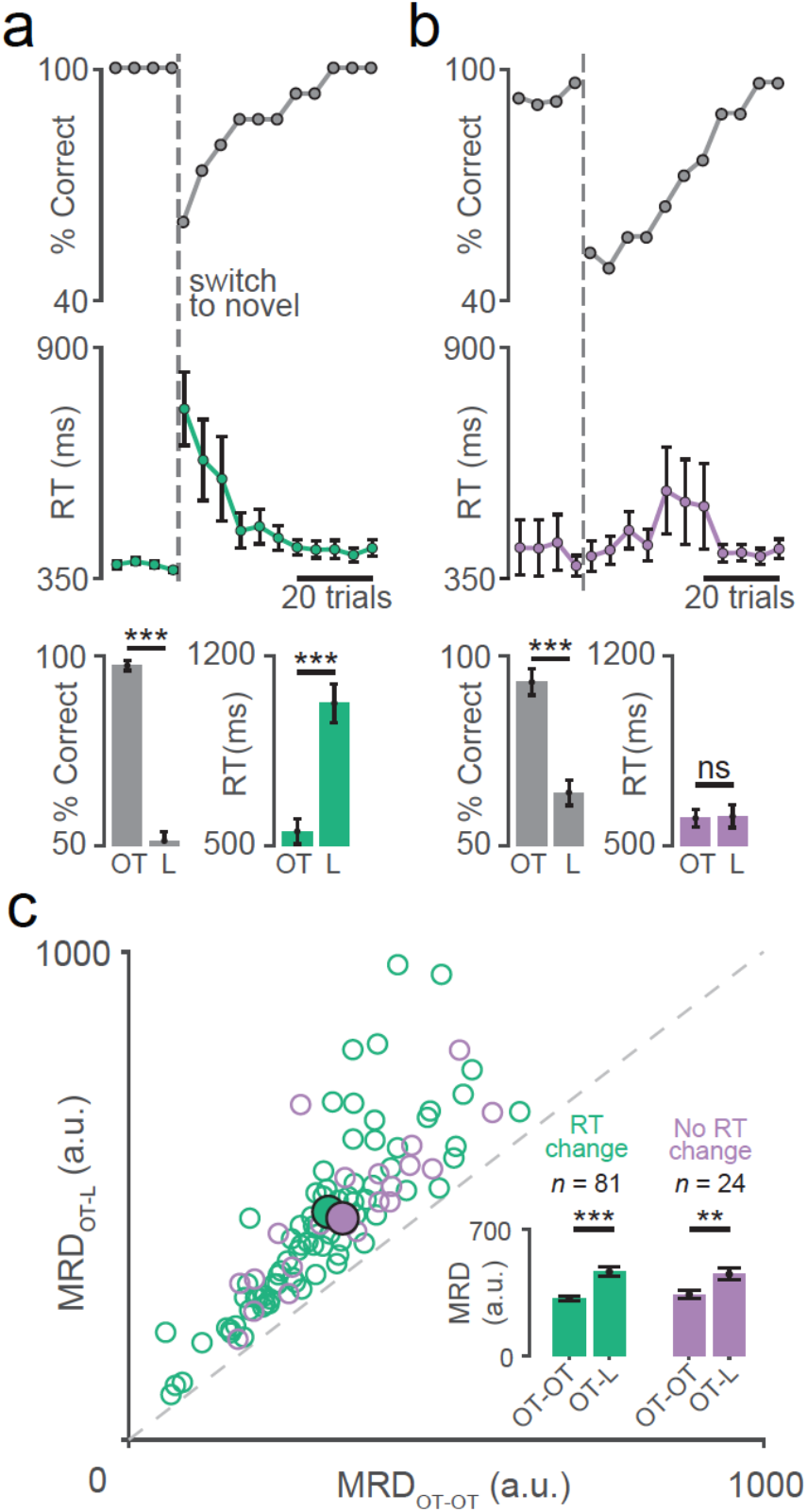
Neural activity changes were independent of changes in reaction time. **a**. Top: Percent of correct trials plotted as a function of trial number relative to the switch to novel visuomotor association. Middle: Reaction times for the same trials. Error bars show the standard error of the mean. Bottom: Mean percent correct (left) and reaction time (right) in the overtrained (OT) and novel learning (L) conditions for all the sessions with changes in reaction time (*** means *P* <0.001, Wilcoxon rank sum test). **b**. Percent correct(top), reaction time (middle), and session averages(bottom) for sessions in which the manual reaction time did not change (n.s means *P* = 0.8151, Wilcoxon rank sum test) after the switch to novel visuomotor association but the performance did (*** means *P* <0.001, Mann-Whitney U-test). Same convention as **a**. **c**. Same data as in **Fig 2c** but separated into sessions with RT change (green; *** means *P* <0.001, Mann-Whitney U-test) and no RT change (violet; ** means *P* <0.001, t-test). condition MRD values indicating that the change in neural activity in the novel condition was significantly different from the activity in the overtrained condition (*P* <0.001 Mann-Whitney U-test; **Fig 2c**).

Although the visuomotor reinforcement learning task was not associated with any change in movement kinematics, we then investigated if forcing a change in kinematics by changing the manipulanda the monkey used to report its decision would be accompanied by a change in the activity of the P-cells, despite the fact that the symbols and their movement association did not change. This experiment began with the monkeys performing the usual overtrained task, starting the task by placing their hands on two bar manipulanda, and reporting their decision by lifting one of their hands. Then, we changed the motor aspects of the task, having the monkeys grab dowel maipulanda (**Fig 4a**) and report their decision by releasing these dowels, which involved a much different movement (**Fig S3**). The kinematics differed significantly between the two versions of the task (**Fig4b**) even though symbols were the same and had the same visuomotor association (**Fig 4c**). The P-cell response did not change when the kinematics changed (example single cell : **Fig. 4d** and population**: Fig 4e** *P* = 0.2330, paired t-test).

**Figure 4:**
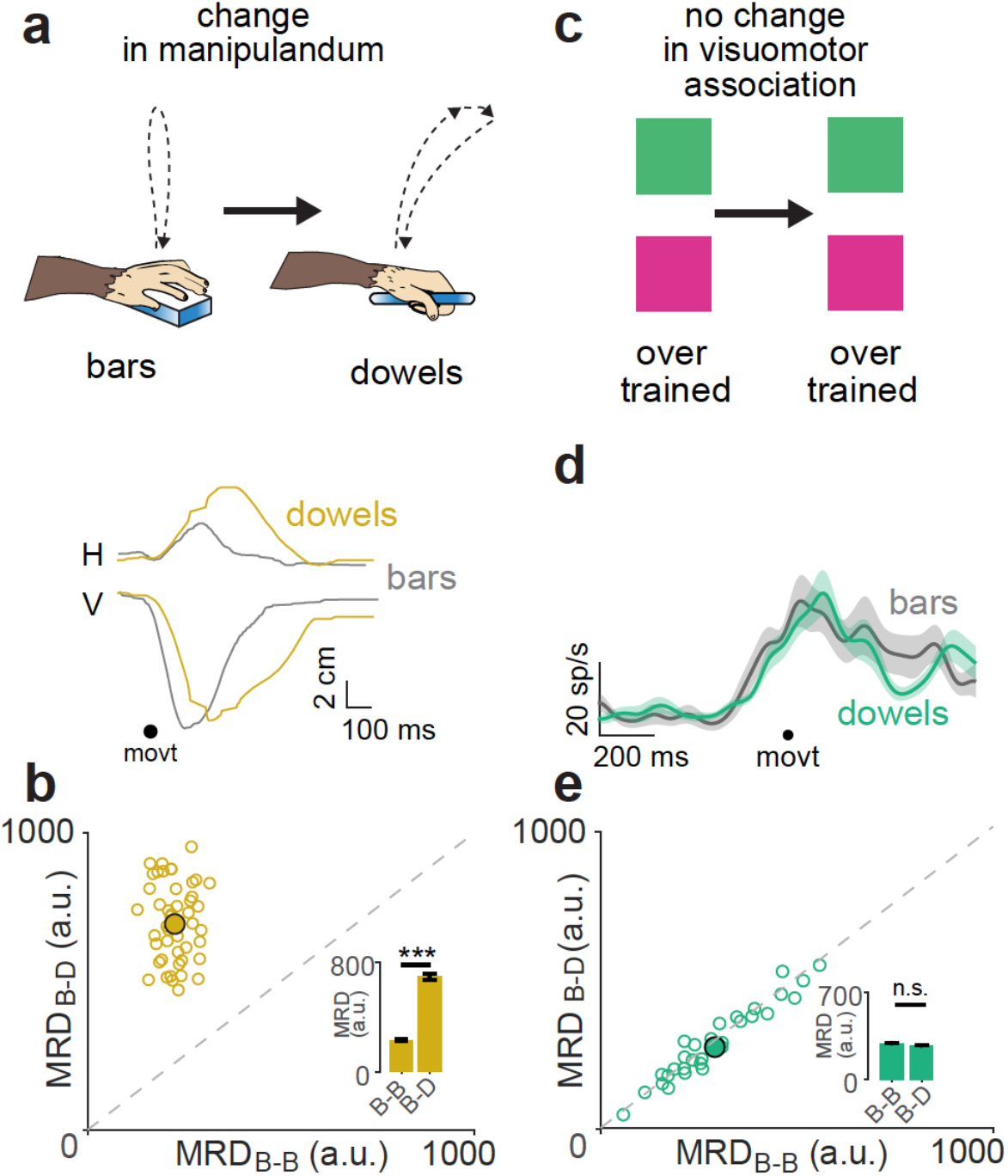
P-cells did not respond to a change in motor kinematics in the absence of a symbol change. a. Top: A cartoon showing different hand movement trajectories with the change in manipulanda from bars (B) to dowels (D). Bottom: Actual movement trajectories decomposed into horizontal (H) and vertical (V) components traces for bars (grey) and dowels (yellow) conditions. b. Same as **Fig 2e** but for hand movement trajectories in bars-dowels condition. *** means *P*<0.001, paired t-test c. Task in which the visuomotor association did not change. d. Same representative neuron from **Fig 2b** when the movement changed but the association did not. e. Same as **Fig 2c** but for neural activity in bars-dowels condition. n.s means *P* = 0.2330, paired t-test

Next, we investigated whether this change in activity profile were merely due to a switch in symbols or whether it were due to the necessity to learn new associations. To do this, we reversed the symbol-hand association once the monkeys had learned the novel association. In this symbol reversal paradigm (**Fig 5a left**), the visual cues that we presented to the monkey remained the same, but the association between the symbols and the hands reversed. That is, the symbol previously cuing a right-hand movement now cued a left hand movement, and the symbol previously cuing a right hand movement now cued a left hand movement. Immediately after the reversal, as expected, the performance of the monkeys dropped below chance level and the monkeys took a longer time to learn the association (**Fig 5a right**). Again, in this condition, the activity profile of the P-cells changed significantly (P < 0.001, t-test; **Fig 5b**) implying that the change in neuronal activity was dependent on the monkey’s having to learn a new association, but was not related to the changes in the symbols per se.

**Figure 5:**
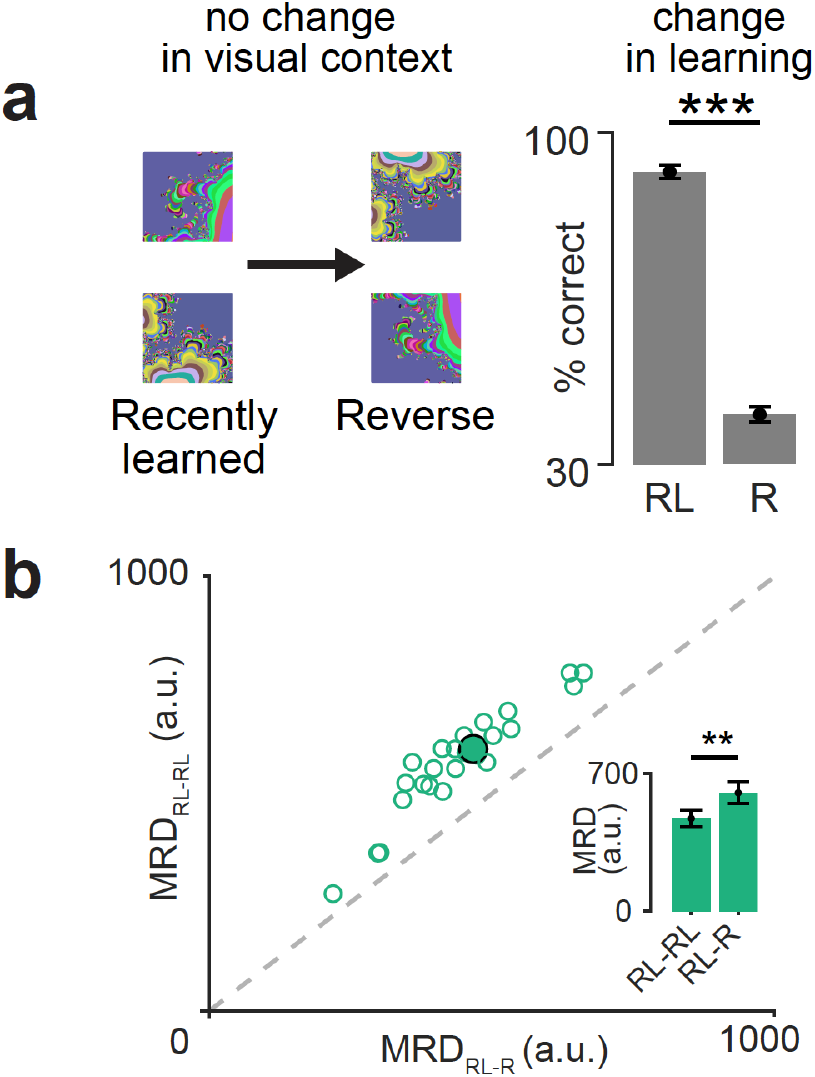
P-cells changed their activity profile in the symbol reversal task. **a**. Left: Changing the visuomotor association of previously learned symbols from recently well-learned (RL) to reversal learning (R) condition but with new learning. Right: Mean behavioral performance in the recently learned (RL) and the reversal (R) conditions. *** means P<0.001, Mann-Whitney U-test. **b**. Scatter plot of MRD for within recently learned condition (RL-RL) vs MRD for across recently learned and reverse condition (RL-R). Same convention as **Fig 2c**. ** means P<0.01, t-test.

### The P-cells changed their activity state as the monkeys learned the task

The transient change in neural activity profile at the beginning of the novel visuomotor association learning (shown in **Fig 2**) propagated through learning, as if the activity state of the cerebellar P-cell network changed. This was similar to previous observations in motor learning paradigms in other studies(Medina and Lisberger, 2008). To show this in a quantitative way, we analyzed the trial over trial changes in neural activity through learning until the monkeys learned the association.

The P-cells changed their activities with learning in three ways: First, 63% the P-cells that showed an increase in firing rate after the symbol switch from the overtrained to novel symbols, continued to increase their firing rate through learning, showing a ‘positive state change’ (for single neuron example, see **Fig 6a**) while the remaining 37% of the neurons eventually returned to the activity, comparable to the overtrained condition, after the initial increase (for single neuron example, see **Fig 6b top panel**), showing no permanent state change. Of the P-cells that showed a reduction in firing rate after the symbol switch from the overtrained to novel symbols, 70% of neurons continued to decrease their firing rate through learning, showing a negative state change (for single neuron example, see **Fig 6c**) while the remaining 30% of the neurons eventually returned to an activity profile comparable to the overtrained condition after the initial decrease (for single neuron example, see **Fig 6b bottom panel**). The neurons did not change the time of their peak activity relative to the movement. Across the population, all three types of neurons were approximately equally distributed (**Fig 6d**).

**Figure 6:**
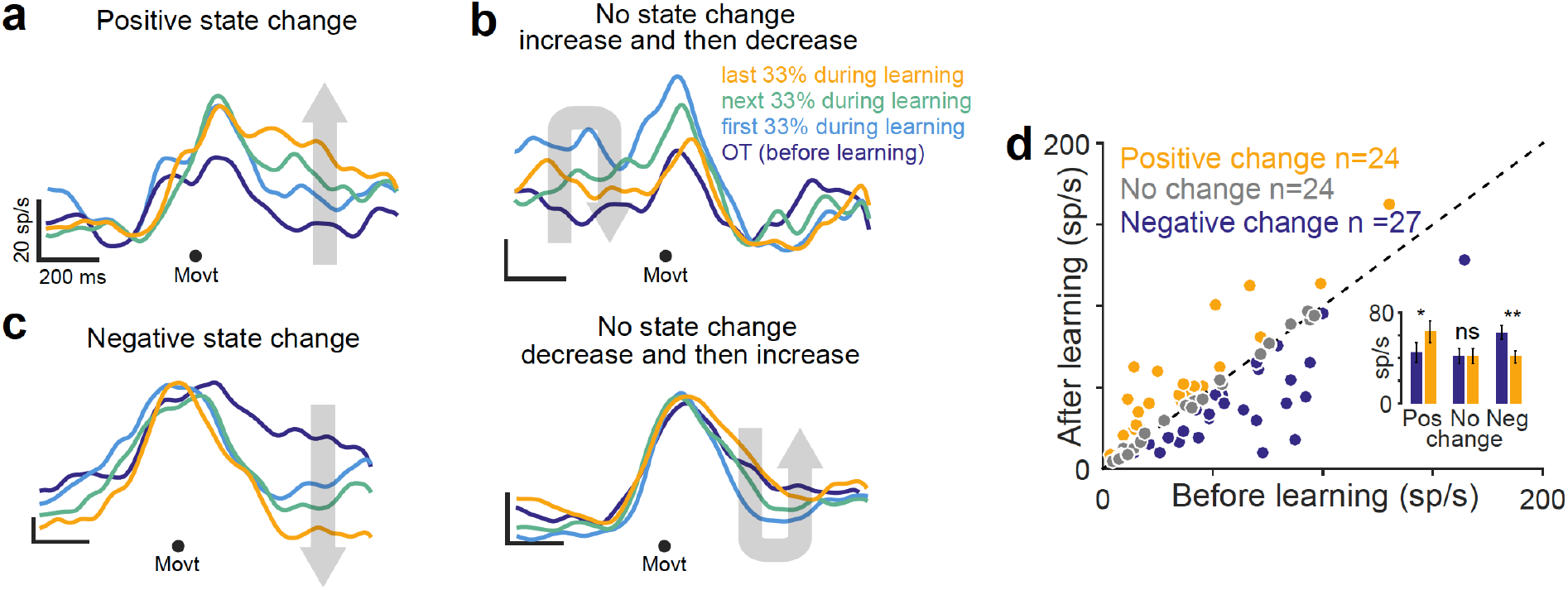
Learning dependent state changes in neural activity. a. Representative neuron whose activity increases at the switch to learning and also increases through learning, showing a positive state change. Activity was synchronized on the movement (dot). b. Top: representative neuron whose activity increases at the switch to learning but decreases through learning, returning to initial state, thus showing no state change. Bottom: representative neuron whose activity decreases at the switch to learning but increases through learning, returning to initial state, thus showing no state change. c. Representative neuron whose activity decreases at the switch to learning and also decreases through learning, showing a negative state change. d. Activity for trials before learning and after learning for all cells. Each dot is a cell. Positive state change neurons lay above the diagonal, negative state change neurons lied below the diagonal. No state change neurons lied on the diagonal. Inset: The mean firing rate for all three classes of neurons before and after learning. * means P<0.05, ** means P<0.01 and n.s. means not significant; t-test.

### Complex spikes did not instruct changes in simple spike neural activity during learning

The conventional model of the cerebellum (Raymond and Medina, 2018) posits that the complex spikes, driven by the climbing fiber input from the inferior olive, provide an error signal that affects the sensitivity of the Purkinje cell simple spikes to the input of the mossy fibers as transmitted by the granule cells. We asked if the complex spike activity (**Fig 7a-b**) induced these learning related changes in simple spike neural activity. We found no relationship between the time of significant modulation of complex spike activity and the time of the epoch of state change for a given P-cell (**Fig 7c-d**). This suggests that there is no obvious relationship between simple spike and complex spike activity during reward-based learning (Larry et al., 2019; Sendhilnathan et al., 2019). Furthermore, if complex spikes were present in the previous trial during learning, the probability that the next trial would be correct was not significantly higher than chance level (**Fig 7e**). This means that complex spikes responses did not affect the simple spike activity or the behavior through an error-based learning mechanism.

**Figure 7:**
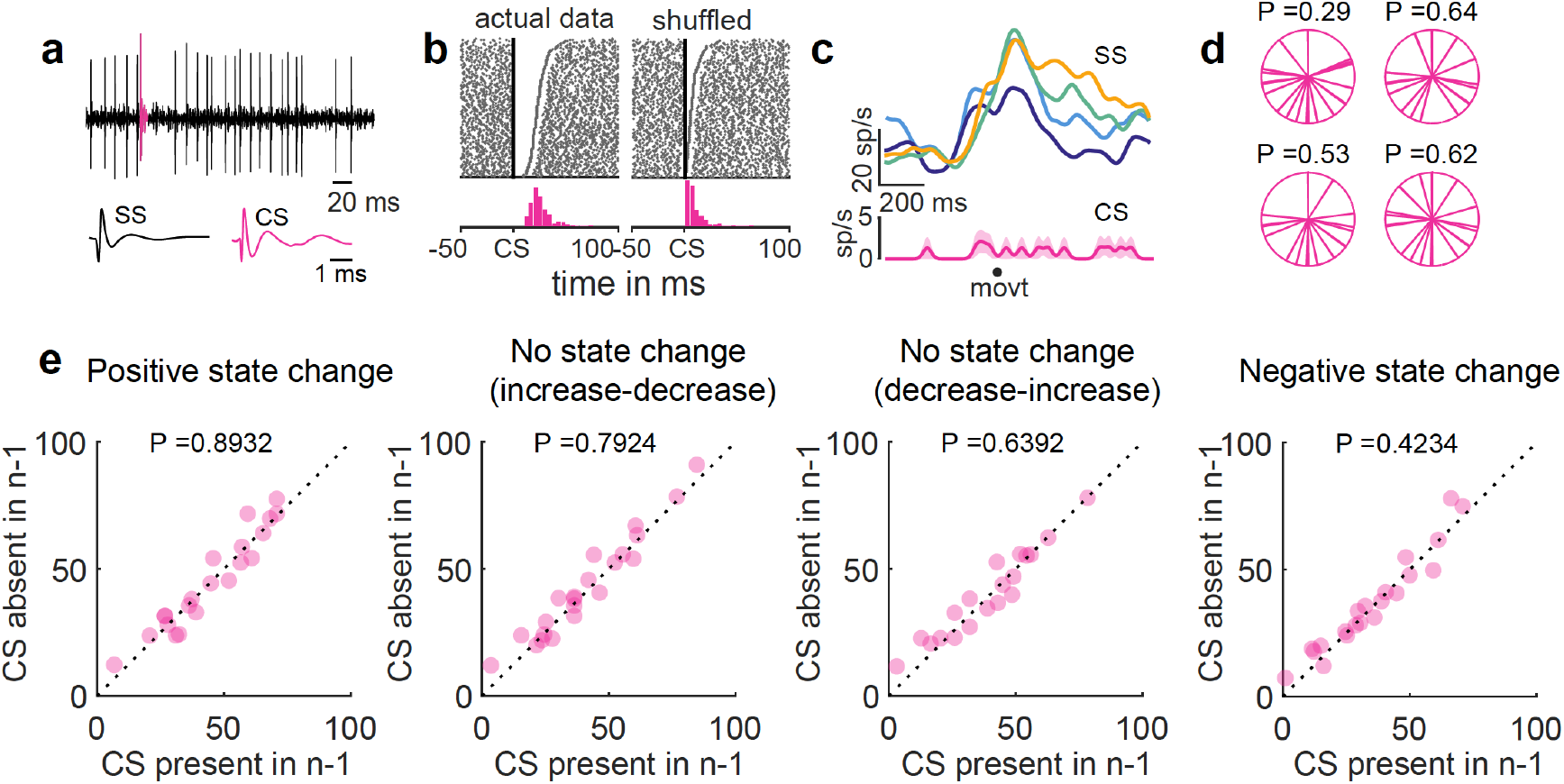
Complex spikes did not instruct simple spikes during reinforcement learning. a. Representative neural recording with simple spikes (gray) and complex spike (pink) b. Complex spike triggered simple spikes for the real data (left) and shuffled (right) c. Top: Same neural response from **Fig 6a** with CS response (bottom). d. Polar plot (of the entire trial period) of time of significant CS modulation during learning relative to the time of state change for each cell. This is shown for positive, negative, no (up-down), no (down-up) state change neurons clockwise from top-left. e. SS activity with CS present in the previous trial (abscissa) vs CS absent in the previous trial (ordinate) for different types of neurons.

## Discussion

We recorded the simple spike and complex spike activities of P-cells in the Crus I and Crus II of the cerebellar hemispheres while monkeys performed a familiar visuomotor association and while they learned a new visuomotor association. We noted several key differences between the neural signatures in these two conditions. First, different neurons showed activity changes in different epochs of the trial at the symbol switch. This change in neural activity was neither due to changes in reaction time or movement kinematics, nor was it due to changes in visual context since the reversal learning followed a similar trend as well. We found that only in the cases that required reinforcement learning of new visuomotor associations did the P-cells change their neural activity to reflect that learning (**Table S1)**.

During learning, induced by the symbol switch, the population of P-cells modulated their response in broadly three different ways. Compared to their neural activity in the familiar task, some neurons kept increasing (positive state) or decreasing (negative state) their firing rate as the animal learned the task and hence their activity after learning the new task was significantly different from the activity in the familiar task. A few other neurons however showed an initial change in activity after the symbol switch, but their activity after-learning returned to a level similar to the activity in the familiar task. Thus, the population of P-cells describes the state of learning without specifying the details of the monkey’s behavior (**Fig 6**).

We previously described a reinforcement error signal (Sendhilnathan et al., 2020) carried by the same P-cells analyzed in this study, that is greatest at the symbol switch and approached zero as the monkeys learned the task, so that the reward activity when the monkeys had learned the novel task resembled that seen in the overtrained task. This was not the case for the state related signal we describe here. While similar to the previously reported reinforcement error signal (Sendhilnathan et al., 2020), the state change shown here only occurred in a limited part of the trial, although the entire population of neurons tiled the entire trial. Although P-cells were sensitive to the outcome on the previous trial, some selective for the previous wrong outcome (wP-cells) while others selective for previous correct outcome (cP-cells) (Sendhilnathan et al., 2020), the nature of the state change was independent of this since we observed similar state change effects for all P-cells (**Fig S5**). Furthermore, although the magnitude of the reinforcement error signal decreased with learning, we found that only some neurons returned their activity (~33%; **Fig 6b**) to their initial state while the rest had significantly different activities between the end of learning and the overtrained conditions (**Fig 6a and c**). Therefore, for these neurons, even though they stop reporting an error signal, their neural activity gets stabilized at a different state, due to learning related changes. This was similar to previous observations in motor learning paradigms in other studies (Medina and Lisberger, 2008).

A hallmark of the previously reported reward based error signal was that it was unlikely to be driven by complex spike activity (Sendhilnathan et al., 2019, 2020). Here too, the concurrent complex spike activity was unlikely to have caused these changes in simple spike activity (**Fig 7**) or could have been the reason why different P-cells had different learning related activities. This is because we did not find any correlation between the time of simple spike activity and the time of complex spike activity. These results are in line with the previous studies which showed a learning dependency on the activity of simple spike, independent of the complex spike in reward based learning (Larry et al., 2019) and motor learning (Avila et al., 2021; Ke et al., 2009; Streng et al., 2018).

Since most of neurons increased the activity before or during the movement, one can argue that these neurons simply encoded the motor command for hand movement, and that the modulation we found depended on the change of the movement kinematic from a familiar to a novel task. However, we can rule out this hypothesis since we did not find any change in the kinematics of the hand movement made by the monkey after the symbol switch (**Fig 2**). Furthermore, when we induced a change in hand movement through a manipulandum switch, despite changes in hand movement kinematics, the P-cell neural activity remained unchanged **(Fig 4**). This strongly suggests that the P-cells did not encode the exact kinematics of the hand movement but rather, goal of the action.

The area of the cerebellum that we have studied lies within a closed loop network involving the prefrontal cortex and the basal ganglia (Bostan et al., 2013). Neurons in these areas are active during visuomotor reinforcement learning, with activity related to reward and movement, although the details of this activity differ from the activity we demonstrated here. For example, in the prefrontal cortex (Asaad et al., 1998) neurons specify the movement that the monkey will make, and learning is manifest by a lengthening of the interval between the neuronal selection of the movement and the actual movement. Conversely, here we show that although the activity of neurons changes during learning, the neurons do not specify the actual movement, nor does the time of their peak activity change relative to the movement. Some prefrontal neurons distinguish between prior reward and prior failure (Histed et al., 2009) but this reward related activity does not seem to change during learning. Both areas show signals with mixed selectivity, the cortex multiplexing selectivity for the symbol and the movement (Rigotti et al., 2013), the cerebellum multiplexing signals for learning error and state. Nonetheless reinforcement learning is impaired by inactivation of prefrontal cortex (Murray et al., 2000) as well as the mid-lateral cerebellum (Sendhilnathan and Goldberg, 2020). How these different aspects of mixed selectivity enable reinforcement learning is a puzzle that remains to be solved.

## Acknowledgments

We thank Glen Duncan for highly creative and wonderful electronic assistance, John Caban and Matthew Hasday for superb machining, Dr. Girma Asfaw, Dr. Moshe Shalev for animal care, Whitney Thomas and Holly Cline for facilitating everything.

## Data and materials availability

Raw data are available online at https://doi.org/10.17632/22n9ps5rzv.1.

## Supplementary Figures

**Figure S1:**
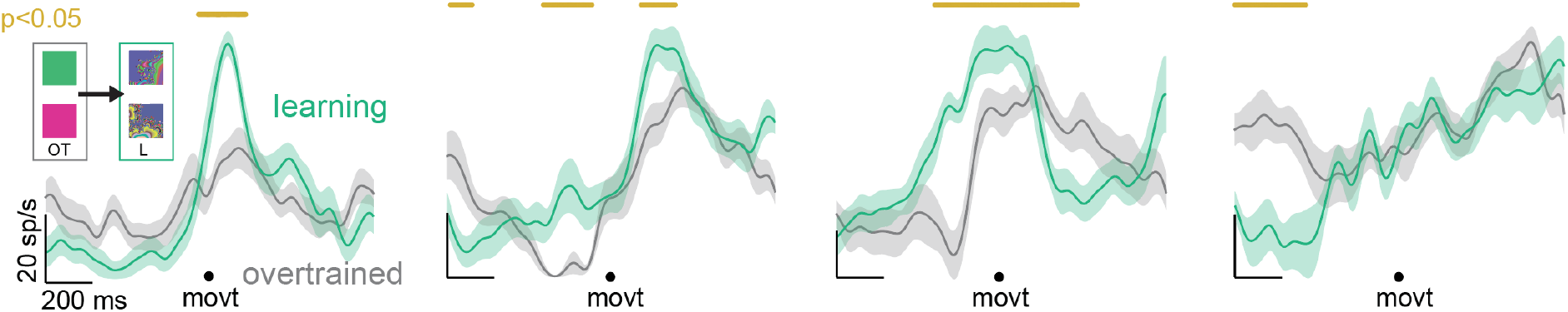
Single neuron examples. Spike density functions of neural activities for four representative neurons during overtrained (gray) and novel learning (green) conditions. The epochs in which these two signals were significantly different (P<0.05), after correcting for multiple comparisons (Benjamini & Hochberg/Yekutieli false discovery rate control procedure), are highlighted with yellow line markers on the top.

**Figure S2:**
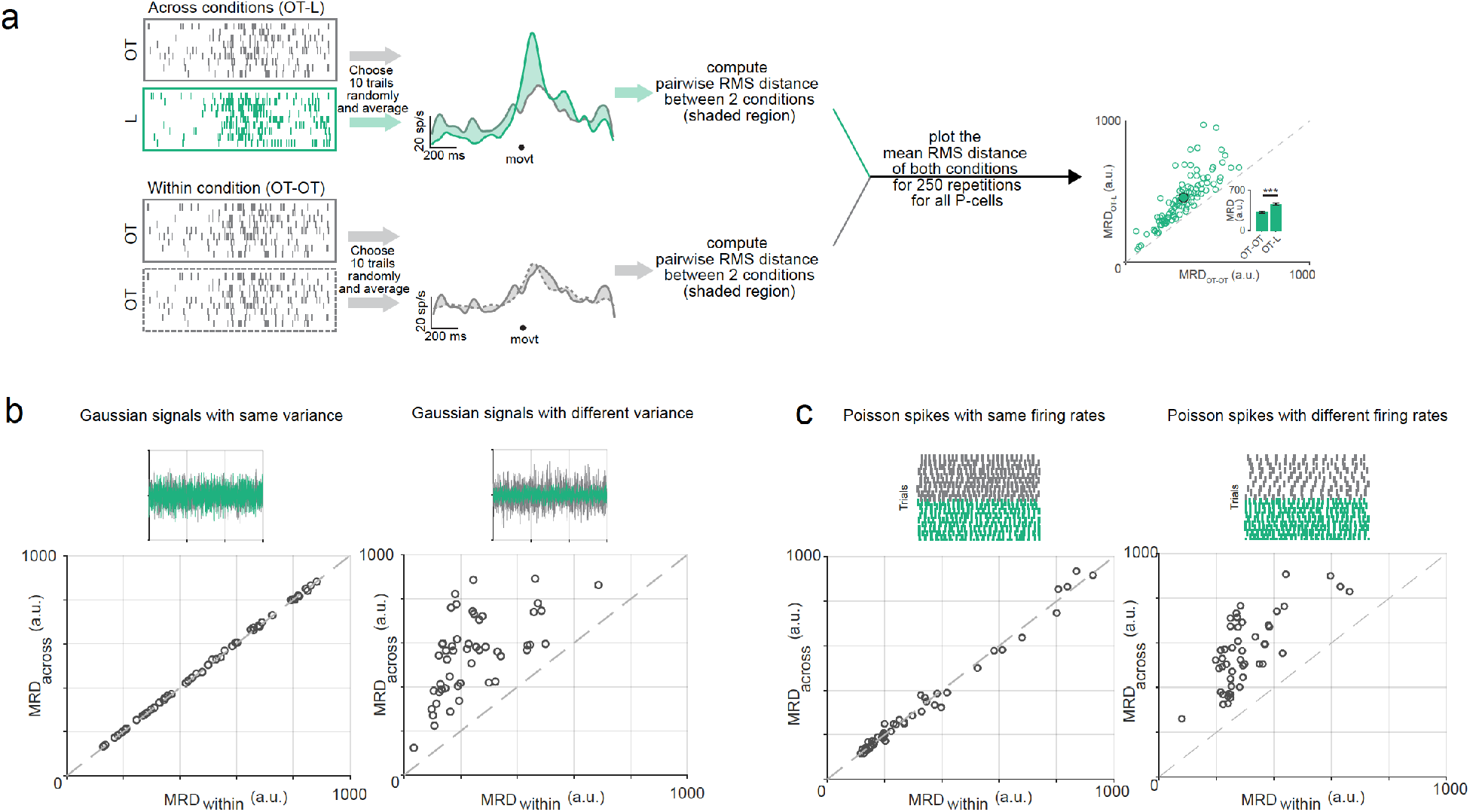
MRD method. **a. Top:** First, we randomly sampled 10 trials each from the last 20 trials in the overtrained condition (grey) and the first 20 trials in the novel condition (green) and calculated the root mean squared (rms) distance between the mean activities. We repeated this process 250 times to obtain a distribution of rms distances that compared the extent of change in across-condition activity profile in the novel condition from the activity profile in the overtrained condition. **Bottom:** To compare this distribution with a control null-distribution, we randomly sampled 10 trials twice without replacement from the overtrained condition and repeated the same analysis to obtain another distribution of rms distances to obtain an estimate of variability of within-condition. A test of statistical significance between the mean of these two distributions would provide an estimate of the change between conditions. **b**. Left: MRD Method applied to 50 pairs of Gaussian signals with 0 mean and same variance. Top panel shows one such pair. Bottom panel shows the Mean RMS Distance (MRD) calculated across group plotted against that calculated within the first group. Right: Same as before, but for 50 pairs of Gaussian signals with 0 mean and different variances. **c**. Left: Same as **b**, but for 50 pairs of mean spike density functions obtained from Poisson spike trains with same mean and variance (λ). Same as b, but for 50 pairs of mean spike density functions obtained from Poisson spike trains with different means and variances (λ).

**Figure S3:**
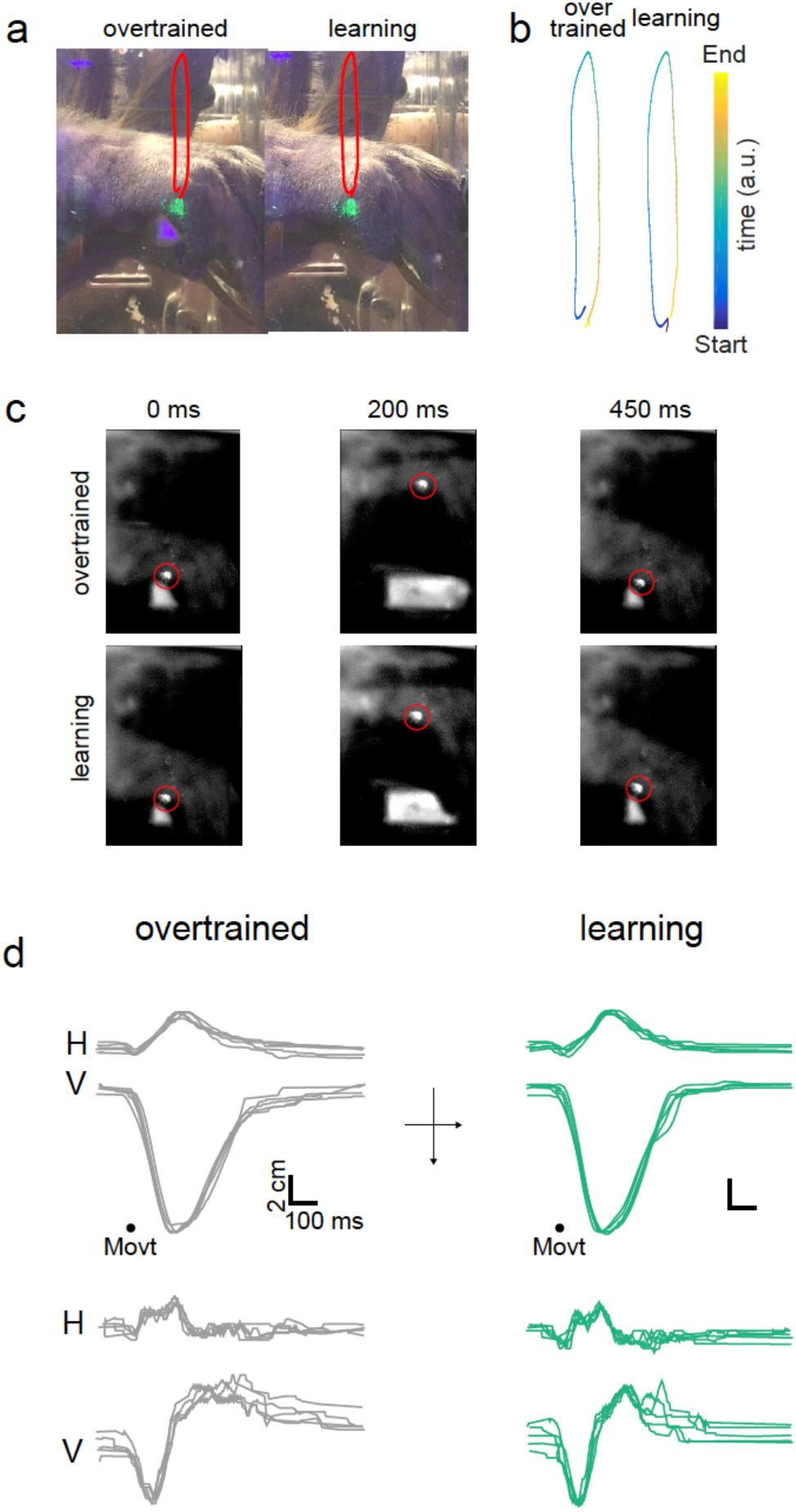
Motor kinematics during symbol switch experiment. **a**. Schematic of the hand movement trajectory as the monkey lifted its hand off the bar in the overtrained (left) and novel conditions (right). The red trace maps the position of the green fluorescent marker in space through the hand movement. **b**. Position of the fluorescent marker in space and time (color indicates the time as in the colorbar inset). **c**. Snapshots from high-frame rate movies showing the monkey’s hand movement trajectory at three time points (0, 400 and 900 ms from the start of movement) for overtrained (top three panels) and novel (bottom three panels) conditions. Red circle marks the fluorescent marker. **d**. Top panel: Horizontal (H) and Vertical (V) hand trajectories for five continuous trials aligned on movement onset for overtrained condition. Bottom panel shows the H and V velocities for the same five trials.

**Figure S4:**
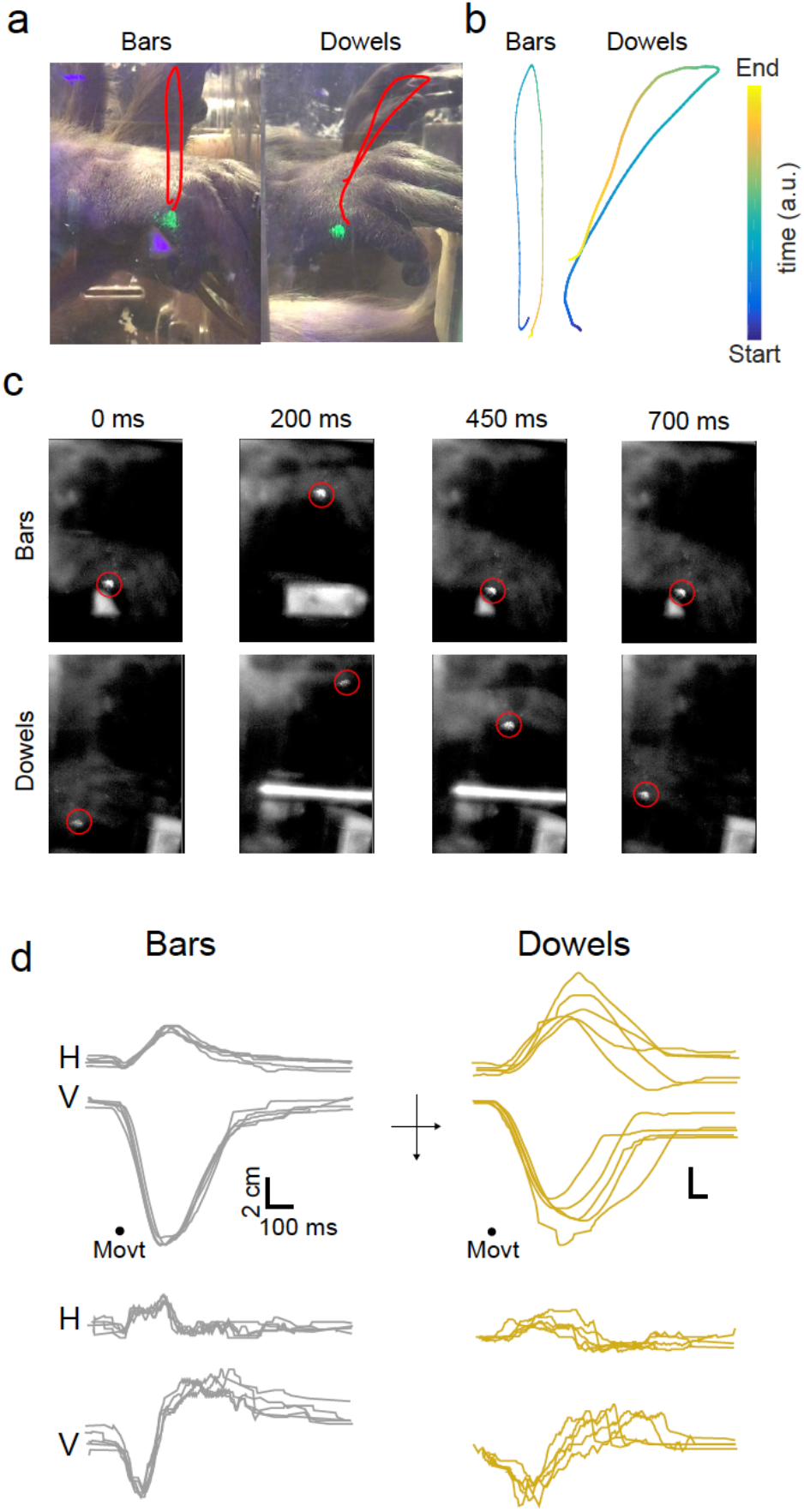
Motor kinematics during manipulanda switch experiment. **A**. Schematic of the hand movement trajectory during bar lift (left) and the dowel release (right) conditions. The red trace maps the position of the green fluorescent marker in space through the hand movement. **B**. Position of the fluorescent marker in space and time (color indicates the time as in the color bar inset). **C**. Snapshots from high-frame rate movies showing the monkey’s hand movement trajectory at four time points (0, 400 and 900 and 1400 ms from the start of movement) for bar lift (top three panels) and dowel release (bottom three panels) conditions. Red circle marks the fluorescent marker. **D**. Top panel: Horizontal (H) and Vertical (V) hand trajectories for five continuous trials with, bar release, aligned on movement onset. Bottom panel shows the H and V velocities for the same five trials.

**Figure S5:**
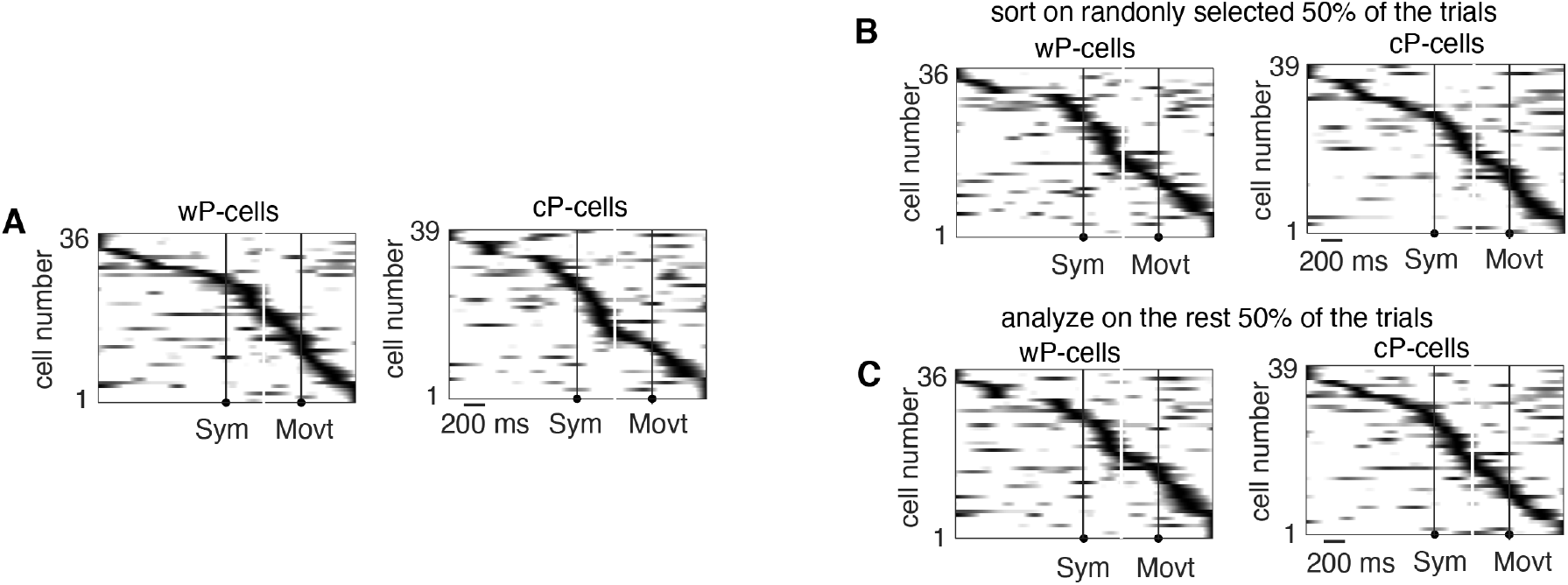
The time of state change tiled the trial period for both cP-cells and wP-cells. A. Heat map of temporally log corrected p values (Benjamini & Hochberg/Yekutieli false discovery rate) from ANOVA among the four waveforms (OT and three phases of learning) for wP-cells (left) and cP-cells (right). B. We sorted the cells by performing the same analysis as in A for randomly chosen 50% of trials (left) and analyzed the cells on the remaining trials (right) as a test of cross validation of the tiling.

**Table S1:** Summary of results.

| Change in<br>parameters<br>Experiment | Change in<br>sensory<br>information | Change in<br>movement | Change in<br>reaction<br>time | Change in<br>reinforcement<br>learning | Change in<br>P-cell<br>activity |
| --- | --- | --- | --- | --- | --- |
| Symbol switch | ✓ |  | ✓ | ✓ | ✓ |
| Symbol reversal |  |  | ✓ | ✓ | ✓ |
| Symbol retrieval | ✓ |  |  |  |  |
| Manipulanda switch |  | ✓ |  |  |  |

## Materials and Methods

We have already described the methods in detail elsewhere (Sendhilnathan et al., 2020). Here we describe it briefly.

### Animal subjects

We used two male adult rhesus monkeys (*Macaca mulatta*), B and S, weighing 10-11 kg each, for the experiments. All experimental protocols were approved by the Animal Care and Use Committees at Columbia University and the New York State Psychiatric Institute, and complied with the guidelines established by the Public Health Service Guide for the Care and Use of Laboratory Animals.

### Task

We used the NIH REX system for behavioral control(Hays et al., 1982). The monkey sat inside a dimly lit recording booth, with its head firmly fixed, in a Crist primate chair 57 mm in front of a back-projection screen upon which visual images were projected by a Hitachi CP-X275 LCD projector controlled by a Dell PC running the NIH VEX graphic system.

### Two-alternative forced-choice discrimination task

The task began with the monkeys grasping two bars, one with each hand, after which a white 1° x 1° square appeared as a trial cue for 800 ms. Then one of a pair of symbols appeared, briefly for 100 ms in some sessions or until the monkey initiated a hand response in other sessions, at the center of gaze. One symbol signaled the monkey to release the left bar and the other to release the right bar. We rewarded the monkeys with a drop of juice for releasing the hand associated with that symbol. We did not punish the monkeys for errors. The monkeys were trained to only release one hand in response to the presented symbol; if they released both hands, the trial was automatically aborted. In the manipulanda change task (**Fig 4**), we began every recording session by presenting the monkeys with the same over-trained symbol pair and bar manipulanda, and after a number of trials, switched the bar manipulanda to dowel manipulanda. The visuomotor association remained the same throughout the task.

## Data collection

### Single unit recording

We introduced glass-coated tungsten electrodes with an impedance of 0.8-1.2 MOhms (FHC) into the left mid-lateral cerebellum (near crus I and II) of monkeys every day that we recorded using a Hitachi microdrive. We passed the raw electrode signal through a FHC Neurocraft head stage, and amplifier, and filtered through a Krohn-Hite filter (bandpass: lowpass 300 Hz to highpass 10 kHz Butterworth), then through a Micro 1401 system, CED electronics. We used the NEI REX-VEX system coupled with Spike2 (CED electronics) for event and neural data acquisition. We verified all recordings off-line to ensure that we had isolated P-cells and that the spike waveforms had not changed throughout the course of each experiment. We identified cerebellar P-cells by the presence of complex spikes online, and offline by the i) spike waveforms, ii) a pause in simple spike after a complex spike, and iii) the simple spike interspike interval distribution(Dijck et al., 2013).

### Hand tracking

We either painted a spot on the monkeys’ right hand with a UV-blacklight reactive paint (Neon Glow Blacklight Body Paint) prior to every session or tattooed the right hand with a spot of UV Black light tattoo ink (Millennium Mom’s Nuclear UV Blacklight Tattoo Ink). We used a 5W DC converted UV black light illuminator to shine light on the spot. Then we used a high speed (250 fps) camera (Edmund Optics), mechanically fixed to the primate chair, to capture a video sequence of the hand movement while the monkeys performed the tasks. We only tracked the monkeys’ right-hand movement because the neurons had similar response with either hand movement. We used the track mate feature(Schindelin et al., 2012; Tinevez et al., 2017) and custom written software in MATLAB to semi-manually track the fluorescent paint spot painted on the monkey’s hand.

## Data Analysis

### MRD method to detect changes in activity pattern

The change in activity for different P-cells occurred at different times in a trial. Given this heterogeneity in its timing, there is no way to predict the time or the magnitude of the activity change, a priori, for a randomly selected P-cell. Therefore, we wanted to use a method that is blind to the time of change, its distribution or the intrinsic property of the neuron and more importantly, makes no a priori predictions about their properties, in such a way that, given the input (for example, activity in OT and activity in N), it classifies activities as same or different across trial conditions. Since the change in activity could occur at any time of the trial for a given neuron, we used the whole trial activity (from before symbol onset through after reward) for this calculation. However, to avoid the trivial possibility where the two activities are different merely due to changes in reaction time latencies, we limited our analyses window to: −400 to 200 ms aligned to symbol onset, and −200 to 600 ms aligned to movement onset for each neuron. The activity aligned to symbol onset and movement onset of the k^th^ trial for each condition (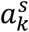 and 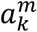 respectively) were concatenated to form single activity vectors for each trial: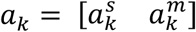.

To quantify the change in activity pattern between two conditions A and B, we computed the mean distance between the two conditions’ activity vectors, where each vector was averaged over 10 randomly drawn trials within each condition.

Clearly, we first identified the condition with least intra-condition variance (suppose condition A). We then compared the activity within the condition with the least intra-condition variance (A) with the activity across both conditions (A and B). To do this, first, we randomly sampled 10 trials each from the last 20 trials in condition A and the first 20 trials in the condition B and calculated the root mean squared (rms) distance between the mean of the sampled 10 concatenated vectors (*a*_k_):

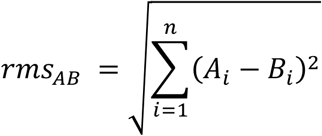

Where 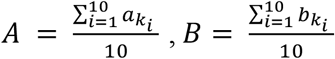 and n is the length of *A* (which is the same as length of *B*), *k*_*i*_ denotes the index of the i^th^ randomly chosen trial.

We repeated this process 250 times to obtain a distribution of rms distances that compared the extent of change in across-condition activity profile between conditions A and B. To compare this distribution with the within-condition activity profile, we randomly sampled 10 trials twice without replacement, from condition A (defined above as the condition with the least intra-condition variance) and repeated the same analysis to obtain another distribution of rms distances.

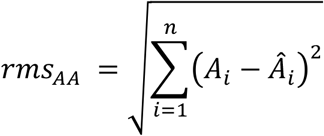

Where 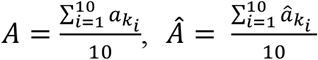 and n is the length of *A* (which is the same as length of Â), *k*_*i*_denotes the index of the i^th^ randomly chosen trial.

We then calculated the means of each of the two rms distributions, called MRD.

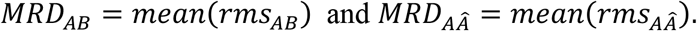

Finally, we repeated this process for each P-cell and performed a statistical test between the population of *MRD*_*AB*_ and *MRD*_*AÂ*_ as shown in see **Fig S2A**.

See **Fig S2B-C** for validity of this method applied to Gaussian distributions and Poisson spike trains. We used this method to compare both neural spike density functions and movement trajectories between different conditions.

### Analysis of state change epoch

We wanted to relate the properties of the state change epoch, to the properties of the delta epoch (Sendhilnathan et al., 2020). Therefore, we followed a similar procedure as that of delta epoch as previously described in (Sendhilnathan et al., 2020) to identify state change so that we can directly compare their properties.

We performed a ANOVA among the simple spike activities in four learning phases: 20 trials in the overtrained, just before the symbol switch, 20 trials just after the symbol switch, 20 trials after 40 trials of symbol switch and 20 trials after learning. For each learning phase, we considered the activity in two epochs: 1) −1400 ms to −400 ms from the symbol onset and 3) −400 ms to 600 ms from the movement onset. We chose this to maximize the sampled trial duration taking the reaction time into account. We chose to extend epoch #1 until 400 ms after the symbol onset and epoch #2 from 400 ms before the movement onset to take into account the long reaction times (800 ms) during early learning trials. After performing an ANOVA in both these epochs, we corrected for multiple comparisons using the Benjamini & Hochberg/Yekutieli false discovery rate control procedure (Q = 0.05). then, we corrected for multiple comparisons across neurons (P values in the state change epoch) using the Bonferroni’s and we cross validated the state change epochs: We sorted the cells on 50% of randomly selected trials and analyzed the data on the held out 50% of the trial and we confirm that the state change epoch’s distribution in time was robust (**Fig S5**).

We defined the start of the state change epoch as the first time-point where the corrected P value became significant and remained significant continuously for the next 150 ms. We defined the end of the state change epoch as the first time-point when the corrected P value became non-significant and stayed non-significant continuously for at least the next 250 ms.

#### Statistics

To check if two independent distributions were significantly different from each other, we first performed a two-sided goodness of fit Lilliefors test, to test for normality, then used an appropriate t-test; or else a non-parametric Mann-Whitney U Test. All error bars and shading in this study, unless stated otherwise, are mean ± SEM.

